# Uncovering functional insights into human pathogenic variants in *CDK19* using Drosophila models

**DOI:** 10.1101/2025.09.10.675453

**Authors:** Karampal S. Grewal, Christopher Tam, Jenny Zhe Liao, Esther M. Verheyen

## Abstract

Heterozygous missense variants in *CDK19* have been found in patients diagnosed with Developmental and epileptic encephalopathy-87 (DEE87) who present with global developmental delay, intellectual disability and other muscular and neurological deficiencies. Two missense variants in CDK19, *Y32H* and *T196A*, were first proposed to be dominant negative in nature based on experiments in a Drosophila model. Subsequently, another group proposed that Y32H is a gain of function based on elevated kinase activity. We present a detailed evaluation of the activity and functionality of these dominant variants in several contexts in fly models of DEE-87 in which endogenous *cdk8*, the fly ortholog of human *CDK8* and *CDK19*, is knocked down while we overexpressed the human genes. Depletion of Drosophila *cdk8* causes thicker muscle myofibrils, fused mitochondria, and climbing defects. The expression of human *CDK19^WT^* in a fly *cdk8-RNAi* background rescues these defects, highlighting functional conservation. To investigate functional differences between the variants, we compared effects of variants to expression of wildtype CDK19. Ubiquitous expression of Y32H can rescue the *cdk8* knockdown phenotype through a gain of function compensatory effect, while T196A is unable to do so through a possible reduction in kinase activity. We found that supplementation of the antioxidant drug, N-acetylcysteine amide (NACA), rescues phenotypes of only the *T196A* adult flies, illustrating a divergence in variant functionality. Our studies in flies allowed us to assay these variants in numerous contexts to gain further insight into their mechanism and obtain translational knowledge to apply back to human health.

## Introduction

Decades of research have established Drosophila as a powerful tool for investigating fundamental biological processes and in understanding human disease mechanisms and functions of rare disease gene variants (1–3). Elegant and diverse assays can be used in flies to probe functional differences between patient variants, which may provide insights into therapeutic interventions. Here we describe our work to understand two patient missense variants in the human CDK19 protein. Cyclin dependent kinase 19 (CDK19) and its paralog, Cyclin dependent kinase 8 (CDK8) are both classified as CDKs that regulate transcription (4,5). They can both bind to their partner cyclin, Cyclin C, and other mediator proteins to form a conserved kinase module that can interact with RNA polymerase II to modulate gene expression, as has been shown across genera (4,6). We recently showed that Cdk8 and CDK19 can regulate mitochondrial dynamics in fly tissues by promoting activity of the fission regulator Dynamin-related protein 1 (Drp1) (7). Furthermore, reduced Cdk8 function can result in build-up of Reactive Oxygen Species (ROS). Aberrant activity or enhanced gene expression of either CDK19 or CDK8 has been linked to diverse ailments in humans, including cancer and developmental syndromes (8–12). A syndrome that is linked to *de novo* heterozygous missense variants of *CDK19* was termed Developmental Epilepsy and Encephalopathy-87 (DEE87) (OMIM 618916) (9,10,13,14). Patients harbouring variants of CDK19 present with global developmental delay, intellectual disability and other neuromuscular deficiencies including ataxia and hypotonia (10).

Drosophila has one ortholog to human CDK19/CDK8 termed Cdk8. *cdk8* mutant flies die at late larval stages (15). Depletion of *cdk8* using RNAi in flies causes muscle defects, seizure susceptibility, and reduced locomotion (7,10). Depletion of *cdk8* can be phenotypically rescued by expression of wild-type human *CDK19* (7,10). This showcases the functional conservation of these orthologues and paves the way to further investigate effects of expressing patient variants in a *cdk8* depleted background as has been done previously to show their pathogenicity (10).

Two *de novo* missense variants of human *CDK19*, *Y32H* and *T196A*, were the first variants of *CDK19* to be characterized (10). The variant Y32H, in which a conserved tyrosine is replaced by a histidine, resides in the Gly-rich loop implicated in ATP binding. In the T196A variant, a threonine is replaced by an alanine at amino acid position 196 which is located within the predicted activation loop implicated in kinase activation and substrate phosphorylation (10,14). The clinical description of the variant expressing patients was accompanied by experiments in Drosophila to gain functional insight into the variants. Using an RNAi-based approach to severely deplete fly *cdk8*, the patient variants *Y32H* and *T196A* were then expressed using the GAL4-UAS system, and various fly phenotypes were then assessed. The strategy of these experiments was to compare phenotypes found when expressing either *Y32H* or *T196A* in a *cdk8* depleted background compared to effects of *CDK19^WT^* (WT) in the same background. The authors tested numerous parameters including viability, lifespan, and sensitivity to mechanical stimulus (10). A theme emerged in their work that pointed towards a possible dominant negative nature of these variants. They proposed this based on their results showcasing that depleting *cdk8* and then expressing either *Y32H* or *T196A* led to effects similar or worse than depletion of *cdk8* alone, indicating that the variants are possibly interfering with the function of the residual Cdk8 in the adult flies.

Another group of researchers aimed to dissect the nature of several *de novo* missense variants using a Zebrafish model (14). In their work, they looked at Y32H and assessed its kinase activity *in vitro* by determining the phospho-status of known targets of CDK19 (14). They observed that Y32H had elevated autophosphorylation and target phosphorylation compared to CDK19^WT^, leading them to postulate that Y32H may be a gain of function missense variant. While they did not assess T196A directly, they suggested that it may be a loss of kinase function variant like another variant (G28R) they characterized (14). In Zebrafish assays they found that both variants caused dominant pathogenic phenotypes, despite having opposite effects on kinase activity.

In this current study, we aimed to further investigate and reconcile the proposed natures of the variants Y32H and T196A by studying effects of these variants on viability, lifespan, muscle function and morphology. Our lab has previously described numerous defects due to depletion of Drosophila *cdk8* in indirect flight muscles, which served as valuable readouts of variant function (7). In the current study we used a genetic background in which *cdk8* was depleted and the patient variant or wildtype *CDK19* genes were expressed. Under these experimental conditions, ubiquitous depletion of *cdk8* yields reduced lifespan, climbing defects, and muscle and mitochondrial morphology defects that can be phenotypically rescued by expression of *CDK19^WT^*. In addition, we show that expression of *Y32H* or *T196A* in this model is pathogenic at 29°C where the activity of GAL4 driving transgene expression is higher, but at 25°C *Y32H* shows no pathogenicity while *T196A* maintains its pathogenicity causing semi-lethality, suggesting that these experimental conditions would allow us to dissect functions in a less pathogenic context. We show that the *cdk8* knockdown climbing defect and muscle mitochondrial morphology can be phenotypically rescued by CDK19^WT^ and Y32H but not T196A. It was previously shown that fly Cdk8 can phosphorylate Drp1 *in vivo* at S616 to induce mitochondrial fission (7). By extension of functional conservation, the same may be said for CDK19. We find evidence that Drp1 phosphorylation at S616 is elevated in the muscle tissue of *Y32H* flies and significantly reduced in *T196A* flies. This is supportive of gain of function and dominant negative hypotheses for the two variants, respectively. Further we show that *T196A* phenotypes can be phenotypically rescued by supplementation of the antioxidant drug NACA which also effectively reduces the ROS level in *T196A* fly adult muscle to control levels. The *Y32H* muscle mitochondrial morphology is unaffected compared to *CDK19^WT^* control group and there is no notable effect to the ROS level upon NACA supplementation.

In summary, we present our model to further dissect the nature of two *de novo* missense variants that may be applied to study other variants that may be linked to DEE87 or other syndromes in humans. We provide evidence that in numerous contexts Y32H acts as a gain of function variant while T196A acts as a dominant negative or inactive kinase. Notably, both of these effects are dominantly pathogenic and associated with similar clinical presentation. Our findings raise the possibility of assessing NACA supplementation in treating the symptoms of human patients and looking at a variant-specific treatment regime in the future. Our studies further highlight the need for diverse functional analyses to dissect variant function in human syndromes.

## Results

### CDK19 variants Y32H and T196A localize to unique regions in the CDK19 protein possibly in close spatial proximity

The amino acid substitutions in CDK19 associated with DEE87 which we have characterized are found to localize to two distinct regions of the protein, namely the ATP binding domain and the activation domain (Fig. 1A). Numerous additional *de novo* missense CDK19 patient variants have been found in these domains (10,13,14). Upon predictive protein modelling and annotation, these residues may be in close proximity with one another in the three dimensional protein structure (Fig. 1B) (16,17), since the region binding ATP needs to contact the activation segment to phosphorylate substrates (18). Patients have one wild-type allele and one dominantly acting variant allele of *CDK19* (Fig. 1C). We have leveraged the GAL4/UAS system in Drosophila to study Drosophila models for DEE87. In these models, an RNAi-based knockdown of endogenous fly *cdk8* is combined with expression of either wild-type human *CDK19* (*CDK19^WT^*) or a *de novo* missense variant (Fig. 1C). These conditions have been used by other groups to induce varying phenotypic severities (7,10). A summary illustration highlights the patient genetic environment with respect to *CDK*19 and the fly model that can be used to study these patient variants. We find that the exogenous CDK19^W*T*^ and the two variant proteins are expressed at comparable levels in adult tissues (Fig. S1A) and that these proteins are abundant in cytoplasmic and nuclear fractions, consistent with our previous work (7). Given that they are expressed at equivalent amounts, we can compare functional differences based on their phenotypes in fly tissues.

**Figure 1.**
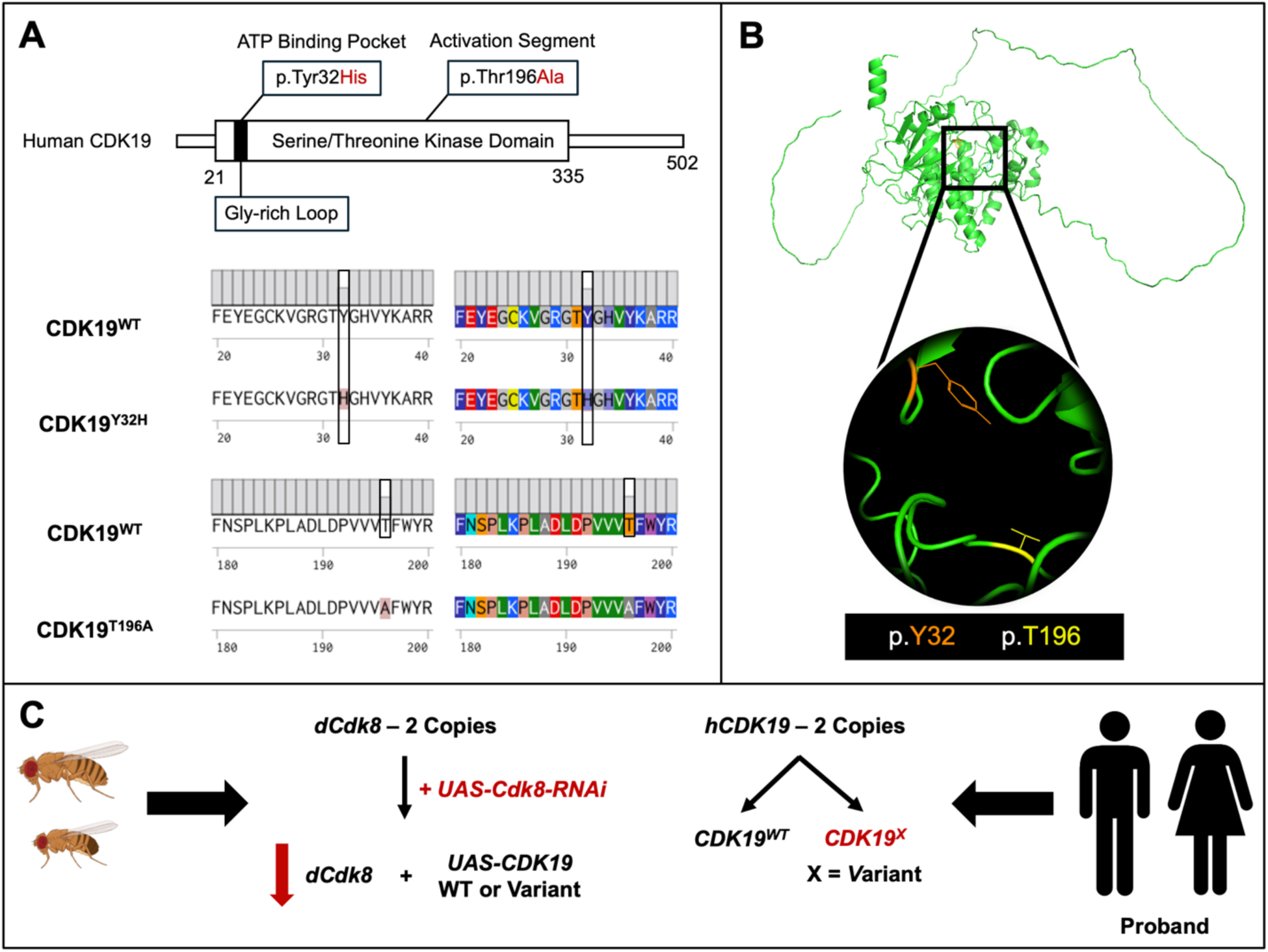
CDK19 variant localization on protein sequence and model. (**A)** Human CDK19 protein sequence cartoon highlighting the amino acid substitutions, p.Tyr32His and p.Thr196Ala, each resulting from a *de novo* missense mutation. Protein sequence alignments of the Y32 and T196 regions for each variant with respect to reference CDK19 using Benchling Molecular Biology Software. **(B)** AlphaFold2 protein model for human CDK19^WT^ (UniProt ID Q9BWU1) with an enlarged field of view of the two amino acid locations: p.Y32 shown in orange and p.T196 shown in yellow. **(C)** Graphical representation of the model used in the paper leveraging the Gal4/UAS system for both expression of a fly *cdk8-RNAi* construct and human *CDK19^X^* cDNA construct (*x = WT, Y32H, T196A*) and the patient genetic environment for context.

### Modulating ectopic expression of *Y32H* and *T196A* in a *cdk8* knockdown background causes variable effects on fly eclosion and lifespan

It has previously been shown that *cdk8* homozygous mutant flies die at early larval stages of development (15) and this effect can be phenotypically rescued upon ubiquitous expression of either fly *cdk8* or human *CDK19* using *actin-Gal4* (10). In contrast, the patient variants *Y32H* and *T196A* are unable to rescue viability in this same background (10). We find that using *daughterless-Gal4* (*da-Gal4*) to perform RNAi-based ubiquitous knockdown of fly *cdk8* at 29°C, yields a reduction in eclosion of viable adult flies (Fig. 2A). Expression of *CDK19^WT^* in this background produces a rescue to control levels (Fig. 2A). Expression of either *Y32H* or *T196A* in this fly *cdk8* depleted background is highly pathogenic and there is a significant reduction in the percent of progeny that eclose as adults compared to the *CDK19^WT^*rescue group (Fig. 2A). We found that at 29°C the percent of pupae eclosed in the Y32H and T196A groups are roughly 3.2% and 0.5% respectively, significantly lower than *cdk8* depletion alone (Fig. S2A). This further reduction in viability in the variant groups compared to the *cdk8-RNAi* group illustrates a dominant effect of these variants on the remaining endogenous fly Cdk8, though it is not clear what causes the pathogenicity.

**Figure 2.**
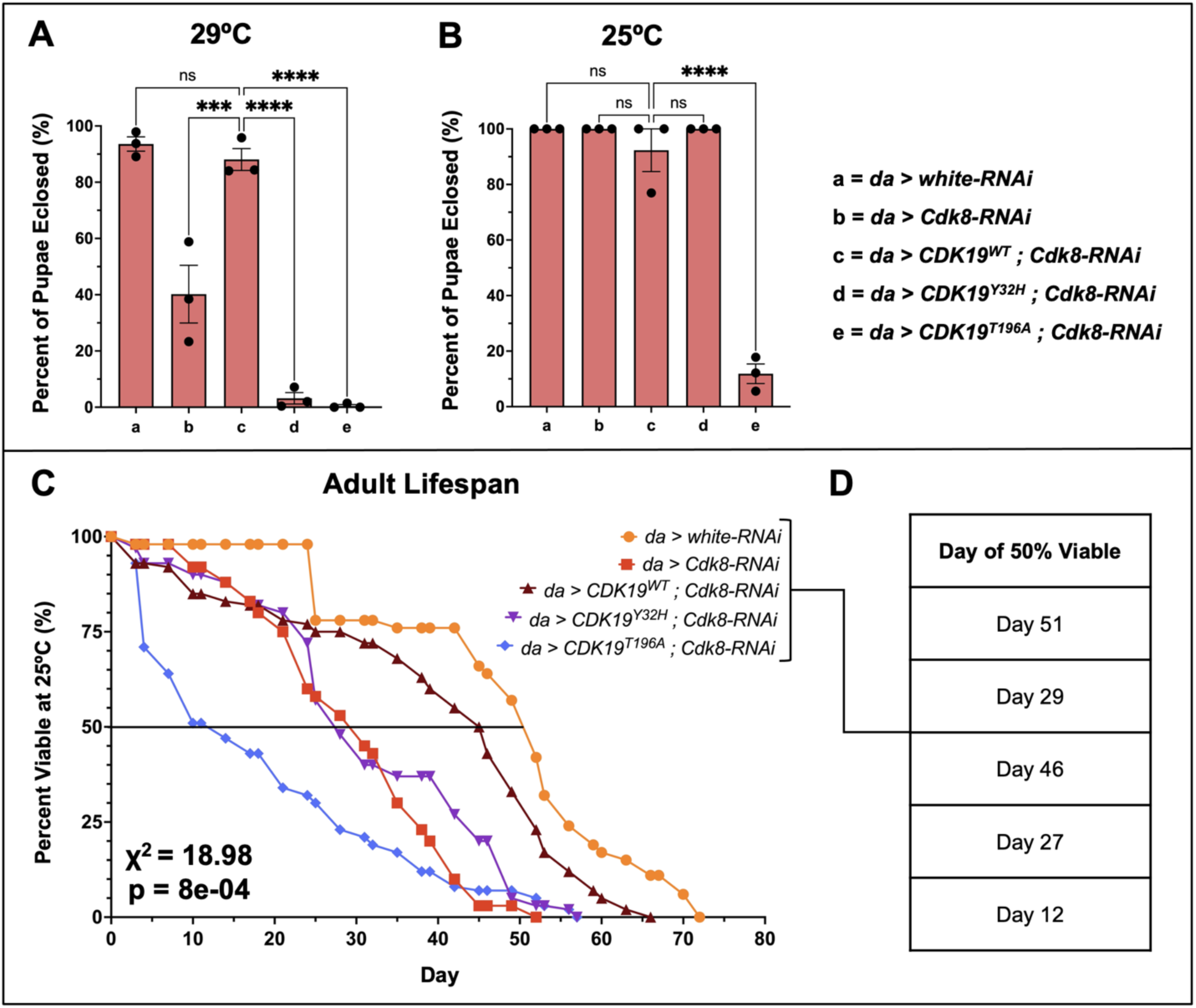
Expression of *Y32H* and *T196A* in a *cdk8* knockdown background causes variable effects to fly eclosion and lifespan. **(A)** Percent of pupae eclosed as adults when ubiquitously expressing the variants in a ubiquitous fly *cdk8* knockdown background at 29°C and **(B)** 25°C (n=3 biological replicates per genotype). Statistical analyses are ordinary one-way ANOVA followed by a Dunnett multiple comparisons test. Results are mean ± s.e.m. (***p < 0.001 ****p < 0.0001). **(C)** Average percent viability of adult male flies for the listed genotypes and **(D)** extrapolated day of 50% viability (n > 45 flies per genotype, n=3 biological replicates). Statistical analyses are a log rank score for survival trends (χ^2^ = 18.98, p = 8.0e-04). Results are average percent viable from n=3 biological replicates.

GAL4 activity is temperature dependent (19), thus raising flies at 25°C instead of 29°C reduces the degree of the *cdk8* knockdown and abrogates the eclosion defect in the *cdk8* knockdown group, resulting in eclosion consistent with control flies (Fig. 2B). Under these experimental conditions *cdk8* was significantly knocked down to roughly 30% of control fly *cdk8* expression levels (Fig. S1B) and all test groups showed a residual *cdk8* expression that is not significantly different from the *cdk8-RNAi* group (Fig. S1C). This less severe knockdown reduces the effect on viability but provides a sensitized background in which to investigate variant functions more fully. Strikingly, we found that expression of *Y32H* in the *cdk8* knockdown background at 25°C did not cause a pathogenic effect on eclosion of adults as was seen at 29°C and there is no significant difference in the percent eclosion between the *cdk8-RNAi* and *CDK19^Y32H^*groups (Fig. S2B). In contrast, expression of *T196A* was dominantly pathogenic in this sensitized background with only roughly 12% of pupae eclosing as adults, significantly lower than the *CDK19^WT^* group (Fig. 2B) and *cdk8* knockdown group (Fig. S2B) respectively.

These findings demonstrate that the variants are functionally distinct, highlighting the need for further research. Thus, we selected the 25°C conditions for our further work to investigate the functional consequences of these two variants, compared to the *CDK19^WT^* and *cdk8* depletion genotypes. This provides two-fold insight, in that we can assess functional conservation or dominant effects of these variants with respect to the wild-type CDK19 and the endogenous fly Cdk8, respectively. Our initial characterizations mirror analyses performed by Chung et al., with the key difference being that in their assays depletion of *cdk8* alone caused 95% lethality, with few escapers, while under our assay conditions, *cdk8* depletion resulted in reduced viability at 29°C but no effect on viability at 25°C. In this context, we could more easily assess distinct effects of the variants.

We next assessed adult fly lifespan at 25°C as a readout for overall fly health by determining how long flies of each genotype lived (Fig. 2C) and extrapolating the time at which 50% of flies had died and 50% still survived (Fig. 2D). Upon carrying out a Kaplan-Meier analysis (log-rank score) to obtain a chi-squared (χ^2^) value for these lifespan data (20), the χ^2^ was 18.98 and from this we obtained a p-value of 8e-04 indicating a significant difference in the lifespan of these groups (Fig. 2C). Under these assay conditions the respective days of 50% viability were 51 days for the *white-RNAi* control group and 29 days for the *cdk8-RNAi* group (Fig. 2D). Expression of *CDK19^WT^* in the *cdk8* depleted background rescues lifespan with 50% viability climbing to around Day 46, while the *Y32H* and *T196A* groups reached 50% viability around Day 27 and Day 12 respectively (Fig. 2D). The *T196A* group showed an early drop off in survival compared to all other groups while *Y32H* behaved like the *cdk8* knockdown alone and the *CDK19^WT^*group performed relatively in line with the control group (Fig. 2C-D). We also found that under these experimental conditions, *Y32H* expression while having no effect to eclosion at 25°C (Fig. 2B), does affect fly lifespan, though *T196A* affects lifespan to a greater severity. We note that there are less viable adult flies from the T196A group compared to the other groups and this severe reduction in the number of *T196A* progeny would imply that the viable flies that successfully eclose are escapers. For this lifespan assessment we had to compensate for this reduction in progenies by amplifying the number of replicate mating vials. To account for this going forward in our work, we selected flies in groups of n = 5 per experimental replicate for subsequent experiments. We observed distinct lethality phenotypes from these *T196A* progenies that were unable to fully develop or eclose. These included elongated L3, a deformed L3-pre-pupa in a pupal case, and some pharate progenies (Fig. S2C).

### *Y32H* can rescue fly *cdk8* depletion climbing defects like wild-type *CDK19* while *T196A* does not

In the lifespan assessment, we noted that the *T196A* group is affected earliest and significantly compared to all other groups, notably the *Y32H* or wild-type *CDK19* groups when comparing the day of 50% viability (Fig. 2C-D). We also noted that there are less viable adult flies from the *T196A* group compared to the other groups. Taking these factors into consideration, we wanted to choose an adult lifecycle timepoint that would allow us to consistently assess phenotypes without running into lethality issues. Therefore, we selected flies at the three-day old stage for further assessments which is a well-used stage across Drosophila literature and also consistent with staging used to assess adult phenotypes caused by *cdk8* depletion (7). These adjustments allowed us to obtain a sustainable number of flies per genotype and perform a greater number of experimental replicates to compensate for the reduced number of *T196A* adults. By controlling for this as the limiting group and normalizing the group assessment size to the *T196A* fly group, our experimental comparisons are more robust as well. With these insights in mind, we recalled that patients harbouring these dominant *de novo* missense variants of *CDK19* present with neuromuscular defects (10) and that these phenotypes are conserved in flies with severely depleted *cdk8* (7,10) we therefore set out to assess adult fly climbing ability as illustrated (Fig. 3A) as a readout for neuromuscular function.

**Figure 3.**
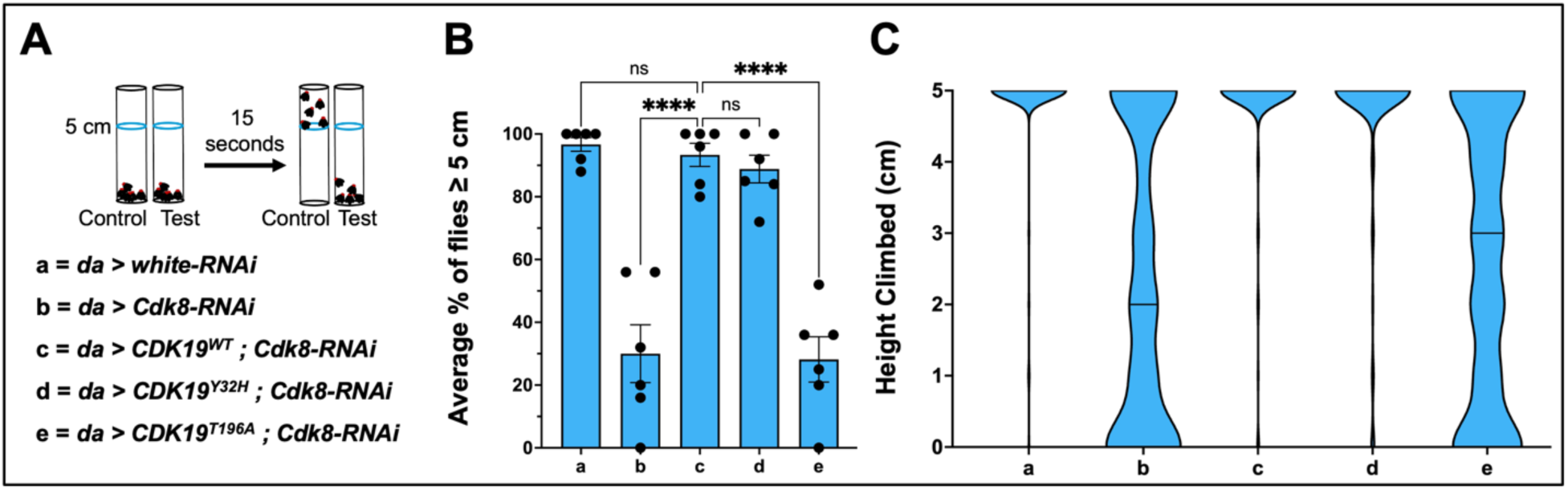
*Y32H* rescues *cdk8* knockdown climbing defects like wild-type *CDK19,* while *T196A* does not. **(A)** Graphical representation of fly climbing assessment for the listed genotypes. **(B)** Average percent of flies that reach ≥ 5 cm (target line) for the indicated genotypes. Each experiment was individually repeated six times (n = 6 biological replicates and n = 30 adult males per genotype). Statistical analyses are ordinary one-way ANOVA followed by a Dunnett multiple comparisons test. Results are shown as bar plots with mean height reached ± s.e.m. (****p < 0.0001, ns no significance). **(C)** Violin plots showcasing climbing distribution as an entire group by plotting the per trial height climbed by each fly for five replicate climbing trials for all independent experimental replicates.

We measured the percentage of flies of a given genotype which could climb to or above the 5 cm line as an endpoint assessment and the height climbed for all tested flies in each individual trial as a group assessment in the set time limit of 15 seconds. Under these experimental conditions, flies with depleted *cdk8* present with significant climbing defects compared to control flies (Fig. S3B) and this is significantly rescued upon expression of *CDK19^WT^* (Fig. 3B). Interestingly, expression of *Y32H* significantly rescued climbing ability compared to the *cdk8-RNAi* group (Fig. S3B) and the climbing ability in the *Y32H* group is not significantly different from the *CDK19^WT^*group (Fig. 3B). This illustrates that both expression of *CDK19^WT^* and *CDK19^Y32H^* can rescue the *cdk8-RNAi* climbing defect to an equivalent degree, indicating an ability of Y32H to compensate for reduction in Cdk8 under these conditions.

One way this result might be rationalized is that this variant has a gain of function as has been proposed previously based on *in vitro* data (14). We showed that expression of *T196A* in a *cdk8* depleted background significantly reduced the number of viable adults indicative of a possible dominant negative effect as previously shown (10) with the number of surviving flies being a fraction of that of the *cdk8-RNAi* or *CDK19^WT^* groups . Here we see that the escaper flies from the *T196A* group have a climbing ability that is not significantly different from the *cdk8* depleted flies (Fig. S3B). It is to be noted that we are dealing with a severely reduced population size in the *T196A* group. Together, these findings illustrate a possible dominant negative effect of T196A on Cdk8 in the flies that were unable to eclose and an inability for T196A to compensate for reduced Cdk8 as CDK19 Y32H and WT can in viable adults.

### *Y32H* and *T196A* flies show similar muscle myofibril thickness and variable muscle mitochondrial phenotypes that correlate to pDrp1^S616^ levels

We see that expression of *Y32H* can phenotypically compensate for reduced endogenous *cdk8* while *T196A* cannot. These findings led us to consider whether there may be a muscular defect underlying both the climbing rescue in the *Y32H* group and impairment in the *T196A* group. We investigated muscle fiber and mitochondria morphologies in the flies expressing the variants. Our rationale was that depletion of endogenous fly *cdk8* has already been shown to lead to muscle fiber thickening and mitochondrial fusion/elongation (7) but no such observations have yet been made for these *de novo* missense patient variants of *CDK19*. We used Drosophila indirect flight muscles (IFMs) to assess both myofibril thickness, sarcomere length, and mitochondrial morphology to assess variant function. Adult IFMs are a well-studied and suitable muscle group to assess these characteristics, they have remarkable similarity to vertebrate cardiac muscle in terms of structure and mode of power output (21,22). IFMs are also known for their use in assessing effects of protein variants on muscle cell function and morphology (21,22).

Under these conditions, ubiquitous *cdk8* depletion yields significantly thicker muscle myofibrils compared to control flies (Fig. S4A) which can be rescued upon expression of wild-type *CDK19* to a myofibril thickness that is not significantly different from the *white-RNAi* control group (Fig. 4A-B, Fig. S4A), as was also shown with muscle-specific knockdown of *cdk8* (7). Both *Y32H* and *T196A* fail to rescue this phenotype with myofibrils significantly thicker than the *CDK19^WT^* group (Fig. 4B) and not significantly different than the *cdk8* knockdown group (Fig. S4A), indicative of a muscle defect upon expression of either variant in the *cdk8* depleted background. We next assessed the sarcomere unit length and observed all groups had consistent sarcomere unit length when using the *CDK19^WT^* group as reference (Fig. 4C). We use the myofibril thickness and sarcomere unit length as proxies for muscle dimensions, a change in myofibril thickness but not sarcomere unit length indicates a net change to fiber thickness.

**Figure 4.**
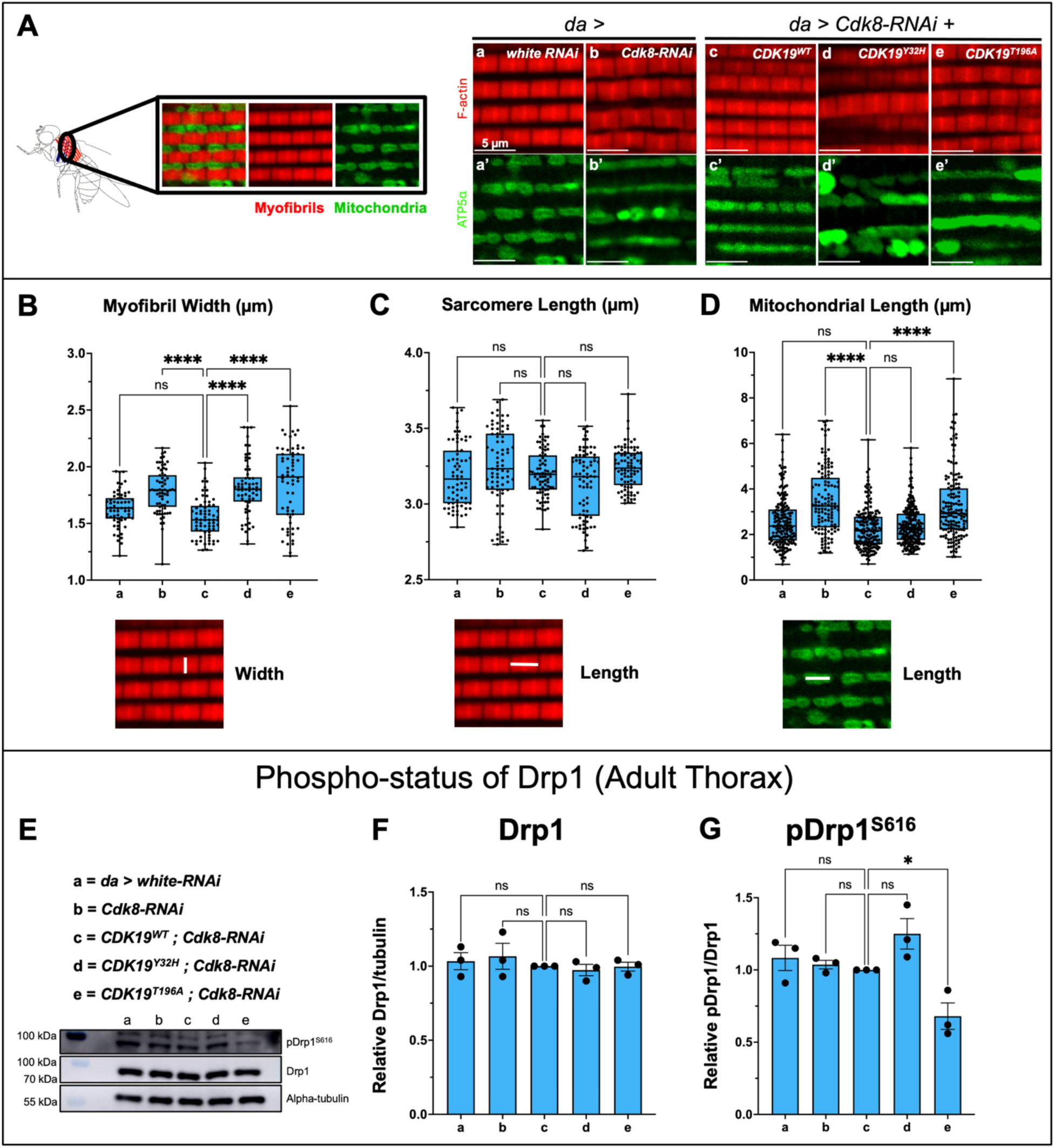
*Y32H* and *T196A* flies show similar muscle myofibril thickness and variable mitochondrial phenotypes that correlate to pDrp1^S616^ levels. (A) Rhodamine-phalloidin staining (F-actin, red) to visualize adult indirect flight muscle for the listed genotypes and ATP5α staining (mitochondria, green) to visualize mitochondrial morphology in the indicated genotypes. Scale bar: 5 µm. **(B)** Quantification of adult fly indirect flight muscle myofibril width. **(C)** Quantification of adult fly indirect flight muscle sarcomere unit length. **(D)** Quantification of adult fly indirect flight muscle mitochondrial length. Quantification of muscle and mitochondria shown as box and whisper plots with solid line at the mean and whiskers for range with 15 muscle regions assessed per genotype. **(E)** Western blot showing the total level of phospho-Drp1^S616^ and Drp1 to assess pDrp1^S616^/Drp1 ratio in the listed genotypes from adult thorax tissue. Densitometry done in Fiji (ImageJ) to show **(F)** total Drp1 protein level and **(G)** phospho-status of Drp1 at position S616 both normalized to alpha-tubulin loading control. Results are presented as bar plots mean intensity ± s.e.m. All muscle tissue is from 3-day old adult males, raised at 25°C, with n = 5 unique muscle tissue per genotype repeated in n=3 biological replicate gels. Statistical analyses are ordinary one-way ANOVA followed by a Dunnett multiple comparisons test. (*p < 0.05, ****p < 0.0001, ns no significance).

We used mitochondrial length as a proxy for morphology as has been done previously when looking at adult IFM mitochondria since mitochondrial shapes are constrained between muscle fibers (7). Under these experimental conditions the *cdk8* depleted group has significantly longer mitochondria than *white-RNAi* control (Fig. S4B) representative of a more fused/elongated phenotype, which can be phenotypically and significantly rescued by wild-type *CDK19* (Fig. 4A,D). Expression of *Y32H* also reduces mitochondrial length compared to the *cdk8* depletion group (Fig. S4B) illustrating a rescue of the *cdk8* depletion mitochondrial morphology defect to a similar degree as *CDK19^WT^* (Fig. 4D). Conversely, *T196A* was unable to do so, and *T196A* mitochondrial lengths were not significantly different from the *cdk8* knockdown group (Fig. S4B) and significantly higher compared to the *CDK19^WT^* group (Fig. 4D). While assessing mitochondrial morphology of the variant expressing groups, we also observed that the *Y32H* group consistently showed more individual mitochondria per assessed area of muscle (Fig. S4C) indicative of a more punctate morphology. Mean mitochondrial length for the *Y32H* group is comparable to the *CDK19^WT^* group and *white-RNAi* control group but there are more individual mitochondria per assessed area on average. We do not see this for the *T196A* group, and this group more resembled the *cdk8* knockdown group (Fig. S4C). These findings showcase that escaper *T196A* flies have phenotypes comparable to the *cdk8* depletion group and T196A is unable to rescue mitochondrial morphology defects caused by *cdk8* depletion while Y32H and CDK19^WT^ can. These T196A escaper flies showcase a possible loss of function nature of this variant and indicate a functional Y32H nature.

The kinase activity of Cdk8 is required for regulation of mitochondrial morphology through phosphorylation of Drp1 at position S616 (7), and given the effects on mitochondria by wildtype CDK19 we propose this function is conserved (7). We assessed the phospho-status of Drp1 (pDrp1^S616^) with respect to total Drp1. This assay provides a two-fold benefit. We can assess the phospho-status of Drp1 to correlate to the mitochondrial morphologies we see in the variant groups, but we can also use this as a functional readout for these two variant kinases by leveraging the conservation between Cdk8/CDK19. For this assay, the adult thorax was chosen for its high muscle to non-muscle tissue ratio. We find that total levels of Drp1 protein relative to tubulin are consistent across all genotypes (Fig. 4E-F, Fig S4D). Under these less severe experimental conditions, *cdk8* knockdown does not cause reduced phospho-Drp1 as the pDrp1:Drp1 ratio is not significantly different from control (Fig. S4E). The pDrp1:Drp1 ratio in the *Y32H* group trends higher, though not significantly. In contrast, the pDrp1:Drp1 ratio is significantly lower in the *T196A* group with respect to the *cdk8* knockdown group (Fig. S4E) and *CDK19^WT^* group (Fig. 4G). Applying the established relationship between the phospho-status of Drp1 and mitochondrial morphology (7) to this result, our data are consistent with the model that Y32H is a gain of function with elevated kinase activity in that the trend in the phospho-Drp1 is higher in this group than the *CDK19^WT^* group, and that T196A is having a dominant negative effect on the remaining endogenous Cdk8 possibly through a loss of kinase activity due to the nature of the variant.

### T196A flies have elevated muscular ROS

We previously showed that *cdk8* depletion correlates with elevated reactive oxygen species (ROS) in neuronal and muscle tissue (7). Given our functional insights and investigations thus far we hypothesise, consistent with previous literature, that Y32H is a gain of function and T196A is acting as a dominant negative. We wanted to assess the effects of expressing these variants in a *cdk8* depleted background on the level of ROS in adult muscle. Our rationale for this assessment is that a gain of function compensation by Y32H in a fly Cdk8 depleted background would likely result in similar or reduced ROS and for T196A we would likely observe an elevation in ROS if this variant were acting on endogenous fly Cdk8 in a dominant negative manner. This assessment can be done using dihydroethidium (DHE) which reacts with superoxide radicals to produce a fluorescent signal that can be detected and subsequently quantified (23).

We observe under our experimental conditions, *cdk8* knockdown flies do not have elevated ROS compared to control *white-RNAi* flies (Fig. S5A), compared to the effects previously shown for targeted muscle-specific knockdown of *cdk8* (7). We observe a trend lower and tightening, though not significant, of the mean fluorescent intensity in the *Y32H* group when comparing both to *cdk8* knockdown (Fig. S5A) and *CDK19^WT^* (Fig. 5A-B). There is a significant increase in the mean DHE fluorescent intensity for the escaper *T196A* flies compared to *cdk8* knockdown (Fig. S5A) and *CDK19^WT^*(Fig. 5A-B). This finding confirmed our hypothesis that the expression of *T196A* leads to elevated ROS in adult muscle and this would be consistent with T196A having a dominant negative effect on the endogenous fly Cdk8.

**Figure 5.**
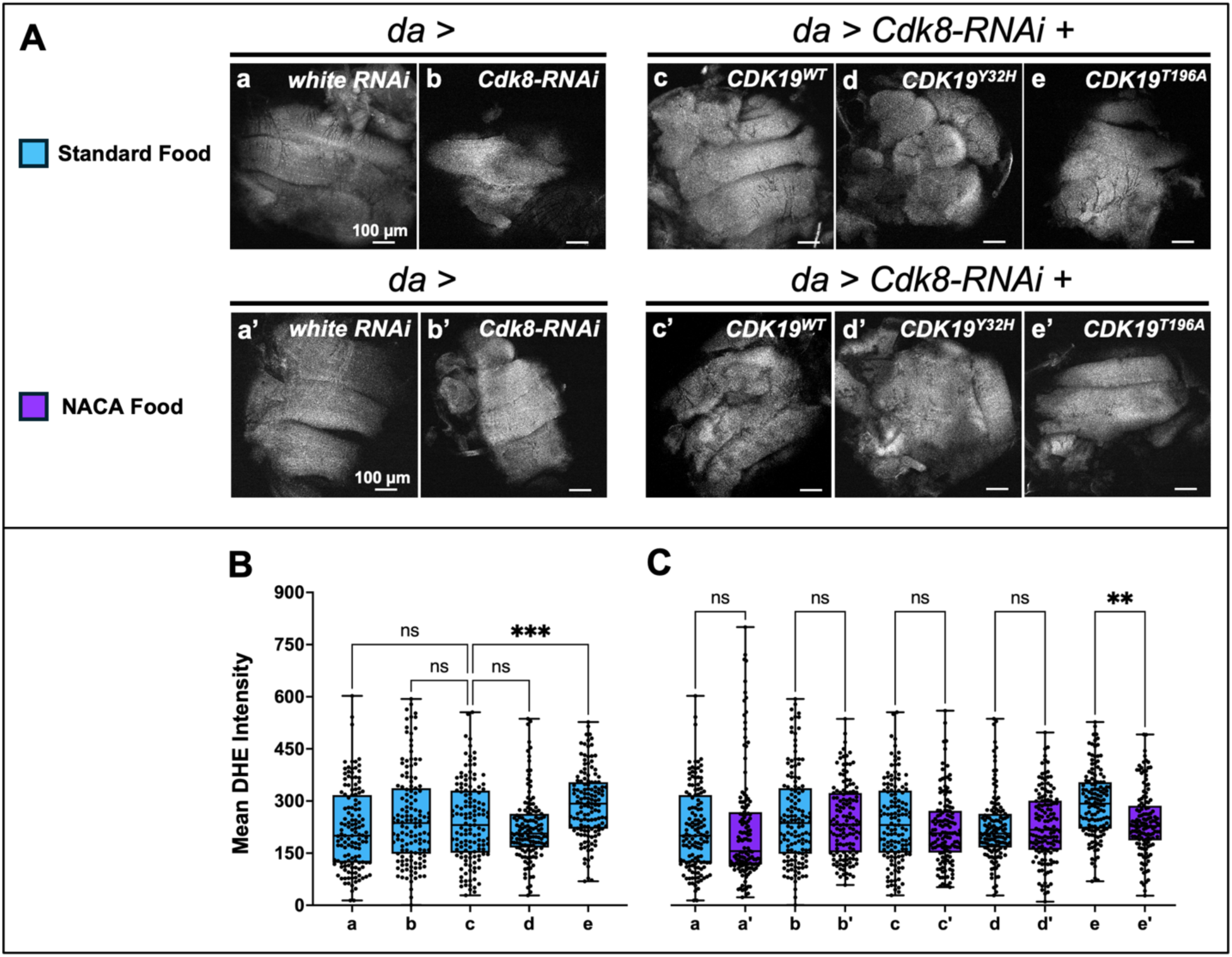
*T196A* adult muscle shows elevated ROS that can be significantly reduced through NACA antioxidant supplementation. **(A)** DHE staining to visualize adult indirect flight muscle (IFM) for the listed genotypes raised on standard food media and NACA supplemented food media. Scale bar: 100 µm. **(B)** Quantification of mean DHE intensity from adult fly IFM from standard food progenies. **(C)** Comparison of standard food and NACA food adult fly IFM DHE intensity. Quantifications shown as box and whisper plots with solid line at the mean and whiskers for range with n = 9 muscle regions assessed per genotype. All muscle tissue is from 3-day old adult males, raised at 25°C, with n=9 muscle across three biological replicates. Statistical analyses are ordinary one-way ANOVA followed by a Dunnett multiple comparisons test. (**p < 0.01, ***p < 0.001, ns no significance).

### Supplementation of the antioxidant NACA can reduce elevated ROS in T196A flies

Given our finding of elevated ROS in *T196A* escaper adult muscle, we wanted to assess if this elevated ROS could be sequestered by supplementation of an antioxidant. Our rationale for this assessment is that antioxidant compounds have previously been used to suppress elevated ROS in flies expressing patient variants of other genes (24). One such antioxidant compound is called NACA, which is a derivatized form of the more than half century old FDA approved drug, N-acetylcysteine (NAC) (25). NACA has a more suitable half-life, improved tissue permeability, and has been shown to be converted into L-cysteine to eventually be integrated into reduced glutathione (GSH) production among other possible routes (25–28). GSH is a chief antioxidant in biological organisms across genera, and we aimed to leverage this NACA/GSH relationship to assess the effect of NACA supplementation on all groups, particularly the escaper *T196A* flies. We supplemented NACA into the fly medium at a concentration of 80 µg/ml, based on a previously determined effective concentration in fly studies (29) and observed that while there was no significant effect to other groups, the escaper *T196A* adult muscle had significantly lower ROS compared to their standard food medium raised counterparts (Fig. 5A,C). The ROS levels in these escaper *T196A* flies raised on NACA is significantly reduced compared to *T196A* flies raised on standard food media to a level that is not significantly different from the other genotype groups (Fig. S5B). Thus, we show that expression of *T196A* is contributing to elevated muscular ROS that NACA can suppress.

### Supplementation of the antioxidant NACA induces a robust shift to punctate mitochondria in adult muscle tissue and climbing ability is improved in T196A flies

It has been well established that mitochondria are the majority contributors to cellular ROS, up to almost 90% (30). We have demonstrated the ability of dietary NACA to suppress elevated ROS in the *T196A* group and further wanted to assess the state of muscle and mitochondria to determine if there is any association with ROS levels and muscle morphology. Our rationale for this assessment is that our lab has previously done a preliminary test on *white-RNAi* and *cdk8-RNAi* IFMs from flies raised with or without NACA supplementation. That experiment showed that NACA supplementation could induce a robust punctate mitochondrial morphology in adult muscle tissue, rescuing the mitochondrial fusion/elongation phenotype resulting from *cdk8* depletion (Fig. S6A-B).

We next investigated the effects of NACA supplementation on the genotypes that we have characterized so far in this study. There is no significant difference in the adult IFM myofibril width from adults raised with or without NACA supplementation (Fig. 6A-B). There is a slight reduction in the escaper *T196A* myofibril width, though not significant. We do find a robust and significant induction of mitochondrial fission resulting in a punctate mitochondrial morphology across all genotype groups (Fig. 6A,C). Most notably, this finding demonstrates that NACA supplementation aids in overcoming the escaper *T196A* mitochondrial fusion/elongation phenotype.

**Figure 6.**
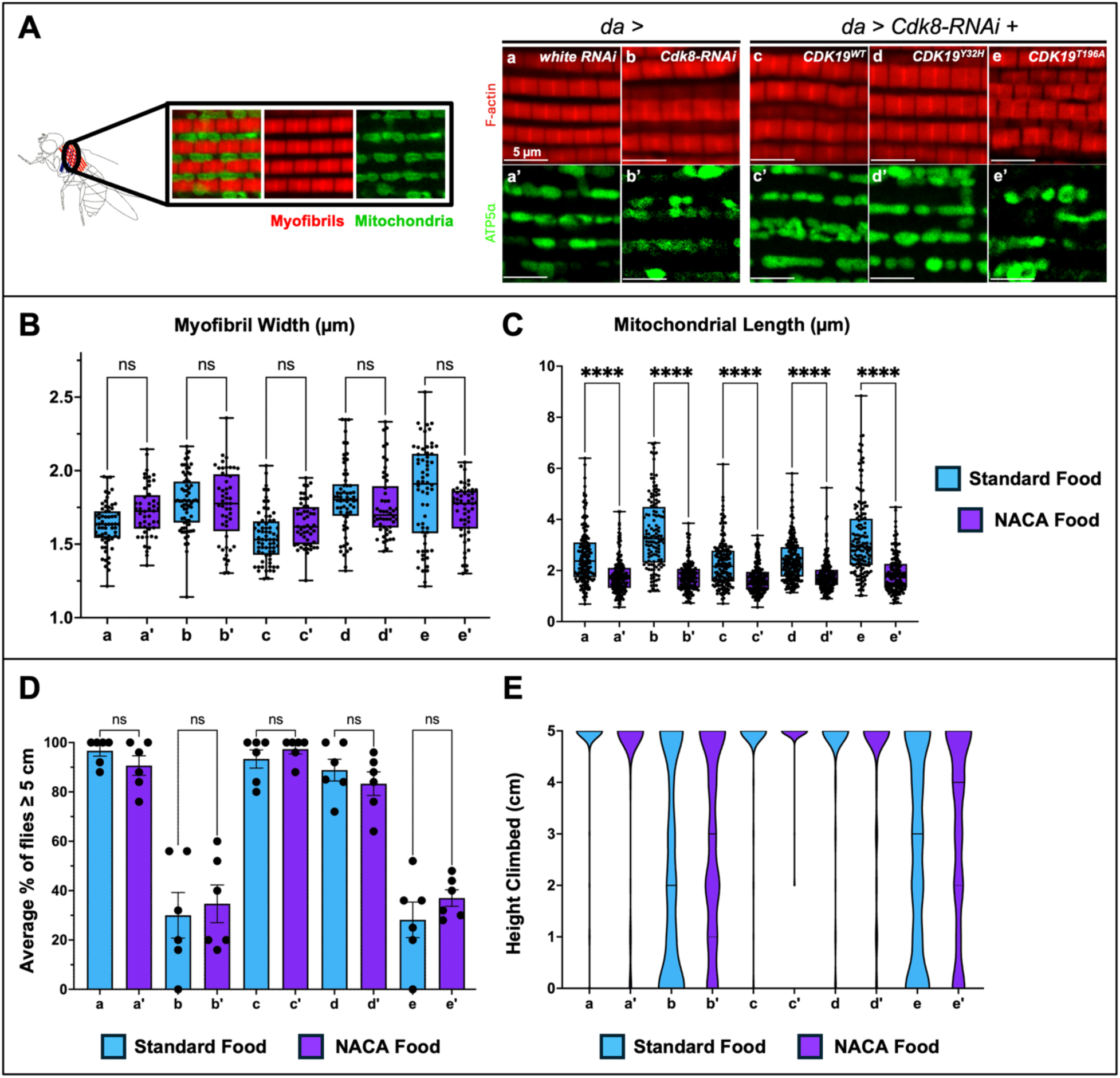
Dietary supplementation of the antioxidant drug, NACA, induces robust mitochondrial fission in adult muscle coupled with climbing improvement in *T196A* flies. (A) Rhodamine-phalloidin staining (F-actin, red) to visualize adult indirect flight muscle for the listed genotypes and ATP5α staining (mitochondria, green) to visualize mitochondrial morphology in the indicated genotypes raised on NACA supplemented food. Scale bar: 5 µm. **(B)** Quantification of adult fly indirect flight muscle myofibril width comparing flies raised with or without NACA supplementation. **(C)** Quantification of adult fly indirect flight muscle mitochondrial length comparing flies with or without NACA supplementation. All muscle tissue is from 3-day old adults, raised at 25°C, with n = 5 muscle per genotype repeated in three biological replicates for a total of 15 unique muscle regions assessed. Data presented as box and whisper plots with solid line at the mean and whiskers for range. **(D)** Average percent of flies that reach ≥ 5 cm (target line) for the indicated genotypes comparing flies raised with or without NACA supplementation. **(E)** Violin plots showcasing climbing distribution as an entire group by plotting the per trial height climbed by each fly for five replicate climbing trials for all independent experimental replicates of flies raised with or without NACA supplementation. Each experiment was individually repeated six times (n = 6 biological replicates and n = 30 adult males per genotype). Statistical analyses are one-way ANOVA followed by a Dunnett multiple comparisons test. (****p < 0.0001, ns no significance).

Following this robust effect of NACA on the mitochondrial phenotypes in adult muscle, we wanted to assess overall neuromuscular health of these flies. We used climbing assessment to compare flies that were raised on standard food vs. NACA food media. Our rationale for this assessment is that the poorest performing adult flies in the climbing assessments harbour elongated/fused mitochondria that are rescued by NACA supplementation. Using the adult IFMs as a readout for muscular health we propose that muscle defects brought on by *cdk8* depletion previously shown (7) or sustained muscle defects upon *T196A* expression in this *cdk8* depleted background are one of the factors creating the locomotion defect we observe in these adult flies. The climbing abilities of these flies were assessed and the *white-RNAi*, *CDK19^WT^*, and *CDK19^Y32H^* groups performed comparably between the NACA groups and standard food groups (Fig. 6D-E), showcasing no effect of NACA supplementation on *Y32H* climbing ability. The *cdk8-RNAi* and *T196A* groups show a trend higher when assessing endpoint height reached with tighter data distribution from the NACA groups notably for *T196A* (Fig. 6D), though not significant. On a per trial basis, however, the *cdk8-RNAi* and *T196A* genotypes both show overall improvement in climbing ability as a group when observing the violin plot distribution for height climbed (Fig. 6E). Interestingly, the *T196A* group performs better than the *cdk8* knockdown group. This indicates that NACA is having a therapeutic effect on the *T196A* flies that may potentially suppress some harmful effects caused by expression of this variant under these experimental conditions. The demonstrated reduction in ROS for the *T196A* group correlates with a more punctate mitochondrial morphology and improved climbing ability as a readout for neuromuscular health and general health of the adult flies.

## Discussion

With the identification of many novel missense variants, including ones of unknown significance, it is important to have robust and diverse assays to study their impacts on protein function. Initial tests of pathogenicity are very powerful to highlight dominant effects of variant proteins, however these do not always reveal underlying activities, depending on experimental conditions. So, while pathogenicity can be established by certain assays, a more in-depth investigation is often needed to identify potential therapeutic measures to avoid ineffective treatments.

Upon initial discovery and characterization of the dominant patient variants of *CDK19^Y32H^*(Y32H) *and CDK19^T196A^* (T196A), a set of Drosophila experiments were carried out to establish pathogenicity (10). These findings showed that these variants were unable to rescue viability when expressed in a *cdk8* deficient background compared to *CDK19^WT^*(WT), highlighting their distinct nature (10). Further, when expressing these variants in a severely depleted *cdk8* background, the phenotypes were largely similar to, or more severe than, *cdk8* depletion on its own (10). The prevailing hypothesis based on these results was that these *de novo* patient variants are loss of function variants presenting as dominant negative. Following this work, another group independently assessed Y32H and other variants using Zebrafish as a model system and they proposed that Y32H may be a gain of function variant with elevated kinase activity while T196A may be a lost/reduced kinase function variant (14). Both assessments used different conditions and model systems. In this study, we aimed to further our understanding of these patient variants by adapting the Drosophila model system previously used to further assess viable adult progeny to investigate various phenotypes and gain functional insight on the nature of these variants. Understanding the biochemical and cellular function of different variants can reveal insights into their clinical features. Of note, our studies confirm the pathogenicity of such variants and reveal that both gain and loss of function can result in similar clinical features.

Here we show under less severe *cdk8* depletion, the *de novo CDK19* patient variant *Y32H* can compensate for reduced fly *cdk8* across numerous phenotypic measures like the reference *CDK19^WT^*can, while *T196A* cannot. We show that under a less severe fly *cdk8* depletion, *Y32H* expression is not pathogenic to the fly host, but *T196A* expression is and there is a significant reduction in the number of viable *T196A* progeny. These limited viable adult progenies are escapers of this genotype, likely representing variability that allows them to complete development and eclose as adults. Further, we reinforce that both variants when expressed under these conditions show pathogenic effects on lifespan compared to control flies; however, upon further dissection we can tease apart subtleties in the effects of these two variants on overall Drosophila health and function.

When we look at adult indirect flight muscle (IFM) to assess effects of the patient variants on muscle cell function relative to the *cdk8-RNAi* and *CDK19^WT^*rescue groups, we show that both variant expressing flies present with thick muscle myofibrils that are not significantly different than *cdk8* depletion alone, highlighting that myofibril thickness is not sensitive to the dominant effects of the variants.

In contrast, when we examined mitochondrial morphology in IFM, we observed distinct effects of the variants, namely the *Y32H* group muscle contains more punctate mitochondria overall and the escaper *T196A* group has more fused/elongated mitochondria overall consistent with the *cdk8* depletion mitochondrial morphology (7). Concomitant to these mitochondrial phenotypes, we provide evidence for elevated pDrp1 levels in the *Y32H* group with respect to the *CDK19^WT^* group and show significantly reduced pDrp1 levels in the *T196A* group compared to both the *cdk8* depletion and *CDK19^WT^* groups. Given the established relationship between Cdk8/CDK19, Drp1 phospho-status, and mitochondrial morphology, we provide evidence from a functional standpoint that Y32H may be phosphorylating Drp1 at position S616 to a greater extent consistent with a gain of function (hypermorphic) kinase nature. Conversely, we show evidence for a significant reduction to Drp1 phosphorylation in the T196A group relative to what we see with *cdk8* knockdown alone, providing further evidence for a dominant negative effect.

Further, we demonstrate that expression of *T196A* is leading to significantly elevated ROS in the adult IFMs. The supplementation of an antioxidant drug, NACA, into the fly food media is sufficient to suppress the elevated ROS in the *T196A* group while having no significant effect on all other genotypes. NACA can be taken up into the cell and utilized for production of GSH (reduced glutathione) (25–28), the chief antioxidant in biological organisms across genera (31,32). NACA has previously been used in Drosophila studies as a therapeutic agent to treat severe neurological phenotypes induced upon expression of patient variants of ACOX1 linked to elevated glial ROS (24).

The supplementation of NACA is also sufficient to induce robust muscle mitochondrial fission in all groups, notably the *T196A* group. These findings coincide with a climbing improvement in *T196A* adult flies and demonstrate that NACA has no significant beneficial or detrimental effect to the *Y32H* variant group. Our Drosophila model allows us to reduce the severity of the *cdk8* depletion to create a viable yet sensitized background to assay the effects of these variants by assessing various phenotypic readouts. It builds on the work that has been done previously and brings further functional insights and investigations into the nature of these variants. This Drosophila model may also, in the future, be adapted to test effects of other patient variants of *CDK19* or its paralog *CDK8*.

In summary, our work reveals further evidence for a gain of function nature for Y32H and dominant negative nature for T196A. Our work also reinforces that both gain of function and loss of function missense variants can lead to human syndromes such as DEE87. Further, we find that there is great potential in assessing the nature of these and other patient variants using fly models and that therapeutic treatments may be prescribed on a variant-specific basis to human patients. We see that NACA supplementation was able to alleviate the severe phenotypes of the *T196A* group while having no detrimental effect to the *Y32H* group, illustrating that the nature of these variants can contribute to unique backgrounds that may be targeted. This allows for the use of a compound such as NACA to target distinct effects caused by *T196A* expression such as abnormally elongated/fused mitochondria and elevated ROS in muscle tissue. These methods may be applied to research into personalized medicinal approaches for DEE87 patients harbouring variants of *CDK19* or other genes linked to various syndromes through a variant specific targeting regime.

## Materials and Methods

### Protein Sequence Alignment and Modelling

FASTA protein sequence of CDK19^WT^ (UniProt ID: Q9BWU1) was aligned to CDK19^Y32H^ and CDK19^T196A^ separately to showcase the amino acid substitutions that arise from single base *de novo* missense mutations (10). To do this the wild-type sequence was manually edited to create separate protein sequences for each variant, Y32H and T196A. The alignment was done in Benchling Molecular Biology Software. To produce a predictive protein model for CDK19, AlphaFold Protein Structure database was used (16,17). The PDB file for this model was input to PyMOL (33) and the protein model was annotated at position 32 and position 196 corresponding to Y and T respectively on the wild-type CDK19 protein.

### Drosophila culture

*Drosophila melanogaster* flies were raised and maintained on a standard cornmeal-molasses medium in accordance with the recipe outlined by BDRC (34) or in standard medium supplemented with 80 µg/ml of N-acetylcysteine amide. Fly stock lines were maintained at 25°C, and crosses were carried out at 25°C and 29°C as indicated. This allowed us to leverage Gal4 temperature sensitivity (19) and modulate degree of expression. The 25°C conditions were suitable to proceed with both variants. The following fly strains were used: *;;da-Gal4* (derived from BL #55850), *;;UAS-Cdk8-RNAi* (BL #35324), *;;UAS-white-RNAi* (BL #33762). The stocks with a BL number are sourced from Bloomington Drosophila Stock Centre (Bloomington, IN, USA). The following stock lines:

*;UAS-CDK19^WT^; UAS-Cdk8-RNAi*

*;UAS-CDK19^Y32H^; UAS-Cdk8-RNAi*

*;UAS-CDK19^T196A^; UAS-Cdk8-RNAi*

were generated in a previous study (10) and generously provided by Dr. Hyunglok Chung (Houston Methodist, Texas, USA).

### Adult Fly Collection

Adult flies were anesthetized using CO_2 (g)_ and collected in styrene vials for use when and where appropriate.

### Eclosion Assessment

Progenies raised at 25°C and 29°C were counted in the pupal stage and following that the number of eclosed adults were counted. The cross for each genotype was carried through three replicate vials and repeated for a total of three biological replicates totalling nine vials per genotype. These data were then quantified and expressed as percent eclosed. This was to assess a readout for viability.

### Lifespan Assessment

Adult flies were maintained at 25°C to assess lifespan. These flies were transferred to fresh food every third day and viability was scored daily on a weekly basis and quantified. This was repeated for a total of three biological replicates and all groups were scored until death. These data were expressed as percent viable per day for each group and a log-rank test was carried out on these data to assess statistical significance of survival trends and to retrieve a χ^2^ value corresponding to a p-value for significance.

### Climbing Assessment

Flies raised at 25°C were subjected to negative geotaxis assays at multiple timepoints: 3-days, 6-days, 10-days and 13-days. The 3-day old assessment was taken as the young-adult timepoint and used for all subsequent assessments. To assess climbing, three-day old adult flies were moved in batches of n = 5 to empty polystyrene vials with another empty vial taped to the top to form a closed cylinder. The bottom vial was marked with dashes at 1 cm increments ranging from 0 cm to 5 cm. The vials containing the flies were gently and simultaneously tapped down and a timer was set for 15 seconds. This trial was repeated for a total of five times These assays were recorded with a Pro 12 MP Digital Camera and the height distribution of each test group was documented and quantified. This test was repeated for a total of five trials per biological replicate.

### qRT-PCR

Drosophila progenies were cleaned in ice-cold 1X PBS and put through an RNA extraction protocol using RNeasy Mini Kits (Qiagen 74134). cDNA for transcript level quantification was produced using OneScript Plus cDNA Synthesis Kit (ABM G236). qRT-PCR was performed using PowerTrack SYBR Master Mix (Applied Biosystems – ThermoFisher – Lot # 29999-452) on a StepOne Real-time PCR System (Applied Biosystems). Primers used:

*rp49* F AGCATACAGGCCCAAGATCG

*rp49* R TGTTGTCGATACCCTTGGGC

*18S F* CTGAGAAACGGCTACCACATC

*18S R* ACCAGACTTGCCCTCCAAT

cdk8 F CATCCGGGTGTTTCTGTCG

*cdk8* R CAGCCCGATGGAACTTAATGAT

To quantify expression level of the target genes of interest, the 2^-ΔΔCt^ method was used (35). The calculations were done manually using *rp49* as the internal reference gene to quantify relative expression level.

### Immunofluorescence staining

Adult thoraces were separated in 1X PBS and fixed in 4% paraformaldehyde (PFA) for 15 minutes at room temperature with rocking. Samples were washed twice for 10 minutes in PBST (0.1% Triton X-100). Tissues were blocked in 5% BSA in PBST for 1 hour at room temperature and then placed in primary antibodies overnight at 4°C. The following primary antibodies were used: mouse anti-ATP5α (1:500, Abcam ab14748). Following primary antibody incubation, tissues were washed with PBST and incubated with the appropriate secondary antibodies: FITC conjugated anti-mouse (1:500, Jackson ImmunoResearch Laboratories) in combination with Rhodamine-Phalloidin prepared in methanol (1:500, ThermoFisher R415) for two hours at room temperature. Samples were mounted on 1 mm spaced glass bridge slides in VectaShield Antifade Mounting Medium (Vector Laboratories, BioLynx) and imaged using a Zeiss LSM880 Airyscan Confocal microscope and processed using Fiji (ImageJ).

### Quantification of mitochondrial morphology

Adult indirect flight muscle images were input to ImageJ and a box of a fixed size was placed to encompass a region of mitochondria. A minimum of 3-5 unique thorax samples were imaged for each genotype repeated for a total of three biological replicates. Lengths of mitochondria were determined manually, and data collected from adult muscle were pooled for box and whisper plot generation and relevant statistics using GraphPad (Prism 10).

### Quantification of muscle fiber width and sarcomere unit length

Adult indirect flight muscle images were input to ImageJ and a box of a fixed size was placed to encompass a region of muscle myofibrils. A minimum of 3-5 thorax samples were imaged for each genotype repeated for a total of three biological replicates. Myofibril widths and sarcomere lengths were determined manually, and data collected from adult muscle were pooled for box and whisper plot generation and relevant statistics using GraphPad (Prism 10).

### ROS Staining and Quantification

Briefly, adult thoraces were isolated in ice-cold PBS and incubated in 30 mM DHE (Cayman Chemical - #12013) for 30 minutes in the dark at room temperature with gentle shaking. Following DHE incubation, tissues were incubated in 8% PFA and washed twice for 5 minutes before mounting on 1 mm spaced glass bridge slides in VectaShield Antifade Mounting Medium (Vector Laboratories, BioLynx). Images were taken within two days using a Zeiss LSM880 Airyscan Confocal microscope. Fluorescent intensity was measured using Fiji (ImageJ). Data collected from adult muscle were pooled for box and whisper plot generation and relevant statistics using GraphPad (Prism 10).

### Statistical analyses

We used GraphPad Prism for statistical analysis and generation of figures. Statistical analysis was done with the default settings of the software (∗ indicates *p* < 0.05, ∗∗ indicates *p* < 0.01, ∗∗∗ indicates *p* < 0.001, ∗∗∗∗ indicates *p* < 0.0001). Violin plots, bar graphs, and box and whisker plots were made using GraphPad Prism 10.

### Web Resources

Benchling Molecular Biology Software

Clustal Omega Alignment Software: https://www.ebi.ac.uk/jdispatcher/msa/clustalo

## Acknowledgements

We thank Dr. Hyunglok Chung (Houston Methodist, Texas, USA) and Dr. Hugo J. Bellen (Baylor College of Medicine, Texas, USA) for the generation of the variant lines used in this study and many productive and engaging discussions. This work was funded by the Canadian Institutes of Health Research to E.M.V. [PJT-495269 (Bridge grant), PJT-189990].

## Abbreviations

ANOVA: Analysis of variation
ATP: Adenosine triphosphate
ATP5α: Adenosine triphosphate synthase subunit 5α
BDRC: Bloomington Drosophila Research Center
BSA: Bovine serum albumin
CDK19: Human Cyclin-dependent kinase 19
CDK8: Human Cyclin-dependent kinase 8
Cdk8: Drosophila Cdk8
cDNA: Complimentary DNA
da-Gal4: Daughterless-Gal4
DEE-87: Human Developmental and Epileptic Encephalopathy Syndrome 87
DHE: Dihydroethidium
Drp1: Dynamin-related protein 1
FITC: Fluorescein isothiocyanate
G28R: Human Cyclin-dependent kinase 19 patient variant G28R
GSH: Glutathione (reduced)
IFM: Drosophila Indirect Flight Muscle
MP: Megapixel
NAC: N-acetylcysteine
NACA: N-acetylcysteine amide
OMIM: Online Mendelian inheritance in man
PBS: Phosphate buffered saline
PBST: Phosphate buffered saline – Tween-20 solution
PCR: Polymerase chain reaction
pDrp1: Phosphorylated Dynamin-related protein 1
PFA: Paraformaldehyde
RNA: Ribonucleic acid
ROS: Reactive oxygen species
s.e.m.: Standard error margin
S616: Amino acid position Serine-616 on Drp1
T196A: Human Cyclin-dependent kinase 19 patient variant T196A
TBST: Tris base – Tween-20 solution
WT: Human Cyclin-dependent kinase 19 wildtype
Y32H: Human Cyclin-dependent kinase 19 patient variant Y32H

---

**Figure S1.**
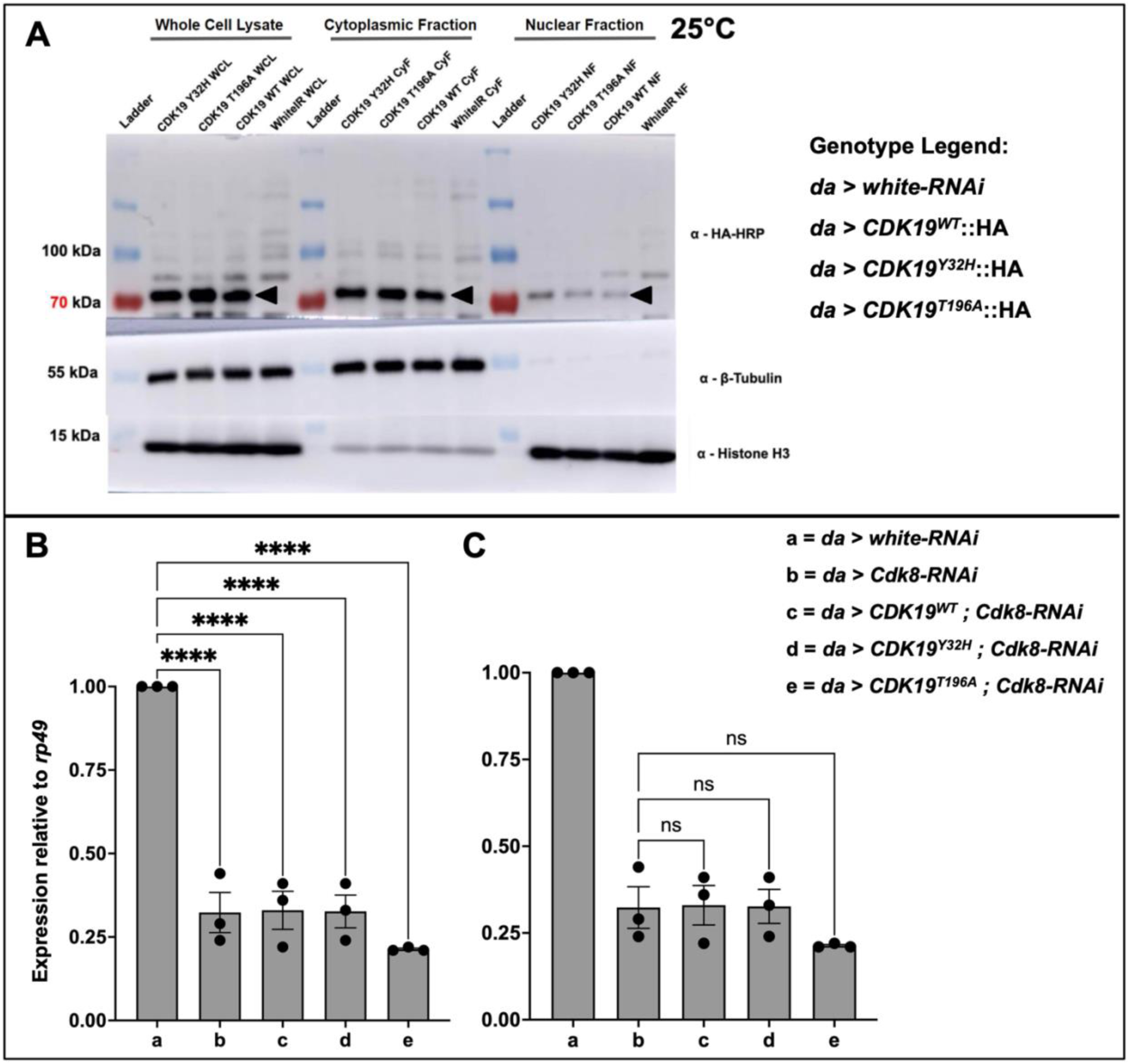
CDK19 proteins have similar levels and localization when expressed ubiquitously in flies and *cdk8* depletion is robust and significant compared to control. (A) Western blot of whole-body thorax lysate fractionation from six whole adult flies using *daughterless-Gal4* to express HA-tagged CDK19 protein constructs at 25°C. CDK19 protein was detected via a 3xHA tag using rat anti-HA-HRP antibody, loading controls were β-tubulin (cytoplasmic) and Histone H3 (nuclear) probed with mouse anti-β-tubulin (abm, G098) and rabbit anti-Histone H3 (CST, #9715) respectively. **(B)** Quantification of expression of *cdk8* (knockdown efficiency) with respect to *rp49* in the listed genotypes compared to control and **(C)** with respect to *cdk8-RNAi* group.

**Figure S2.**
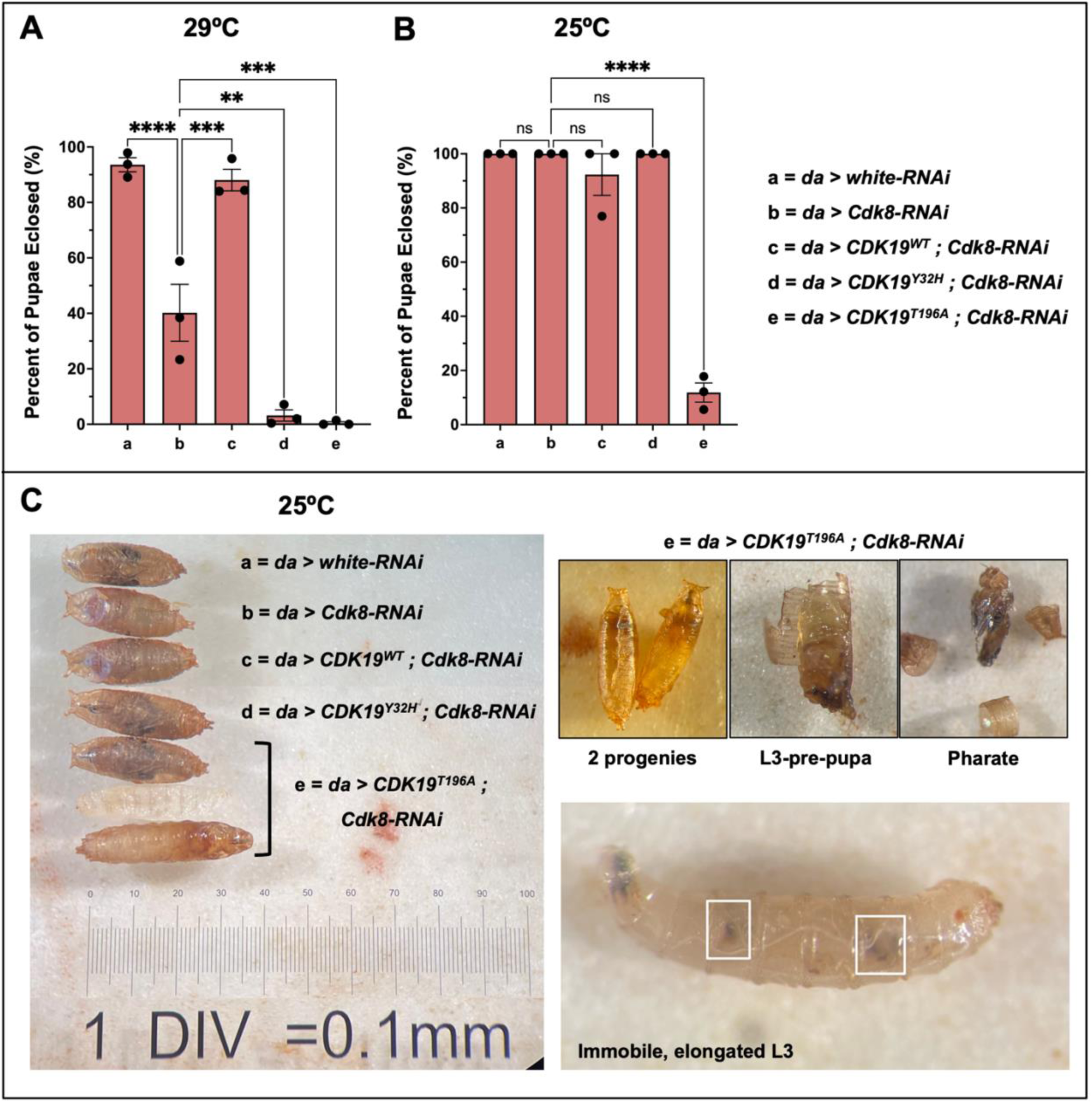
Expression of *Y32H* is pathogenic at 29°C but not at 25°C while *T196A* is pathogenic at both temperatures compared to *cdk8* knockdown alone. (A) Percent of pupae eclosed as adults when ubiquitously expressing the variants in a ubiquitous fly *cdk8* knockdown background at 29°C and **(B)** 25°C (n=3 biological replicates per genotype). **(C)** Pupae of the listed genotypes compared to various progeny obtained from the *T196A* mating vial including elongated L3, undeveloped pupae, and pharate progenies. Statistical analyses are ordinary one-way ANOVA followed by a Dunnett multiple comparisons test. Results are mean ± s.e.m. (**p < 0.01 ***p < 0.001 ****p < 0.0001).

**Figure S3.**
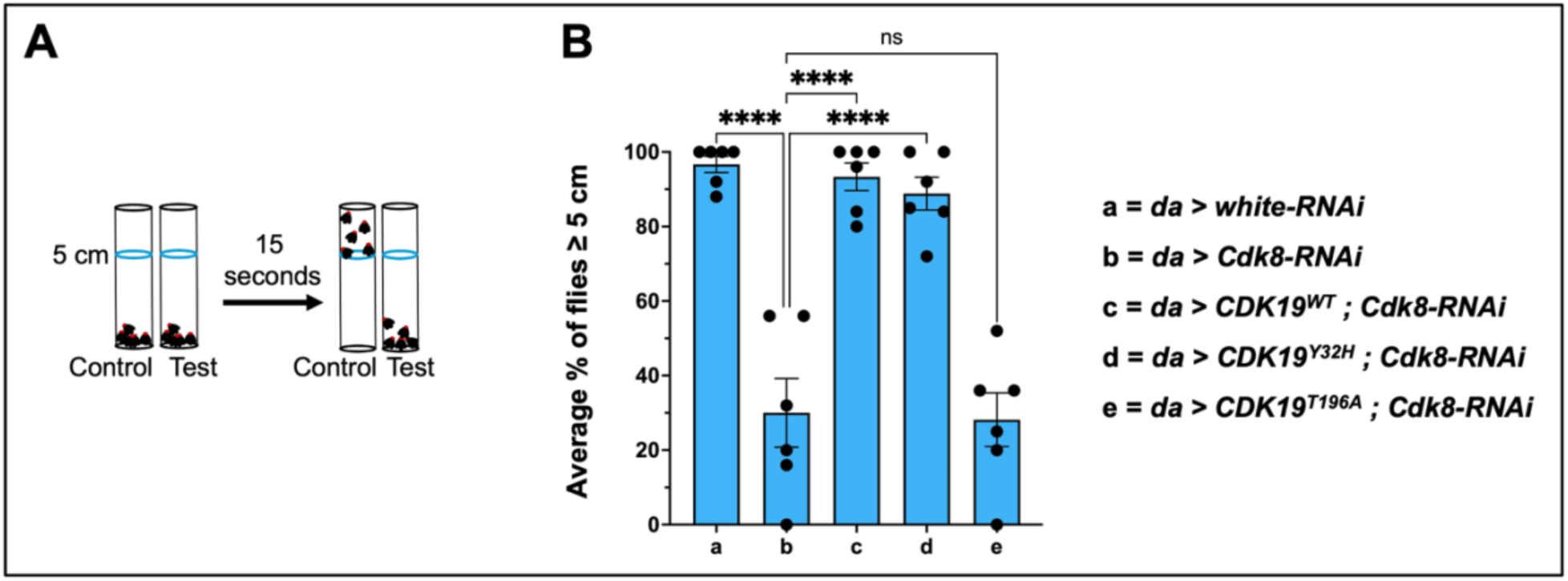
Expression of *CDK19^WT^, CDK19^Y32H^* both significantly rescue *cdk8* while T196A escaper flies are not significantly different in their climbing ability. (A) Graphical representation of fly climbing assessment for the listed genotypes. **(B)** Average percent of flies that reach ≥ 5 cm (target line) for the indicated genotypes. Each experiment was individually repeated six times (n = 6 biological replicates and n = 30 adult males per genotype). Statistical analyses are ordinary one-way ANOVA followed by a Dunnett multiple comparisons test. Results are shown as bar plots with mean height reached ± s.e.m. (****p < 0.0001, ns no significance).

**Figure S4.**
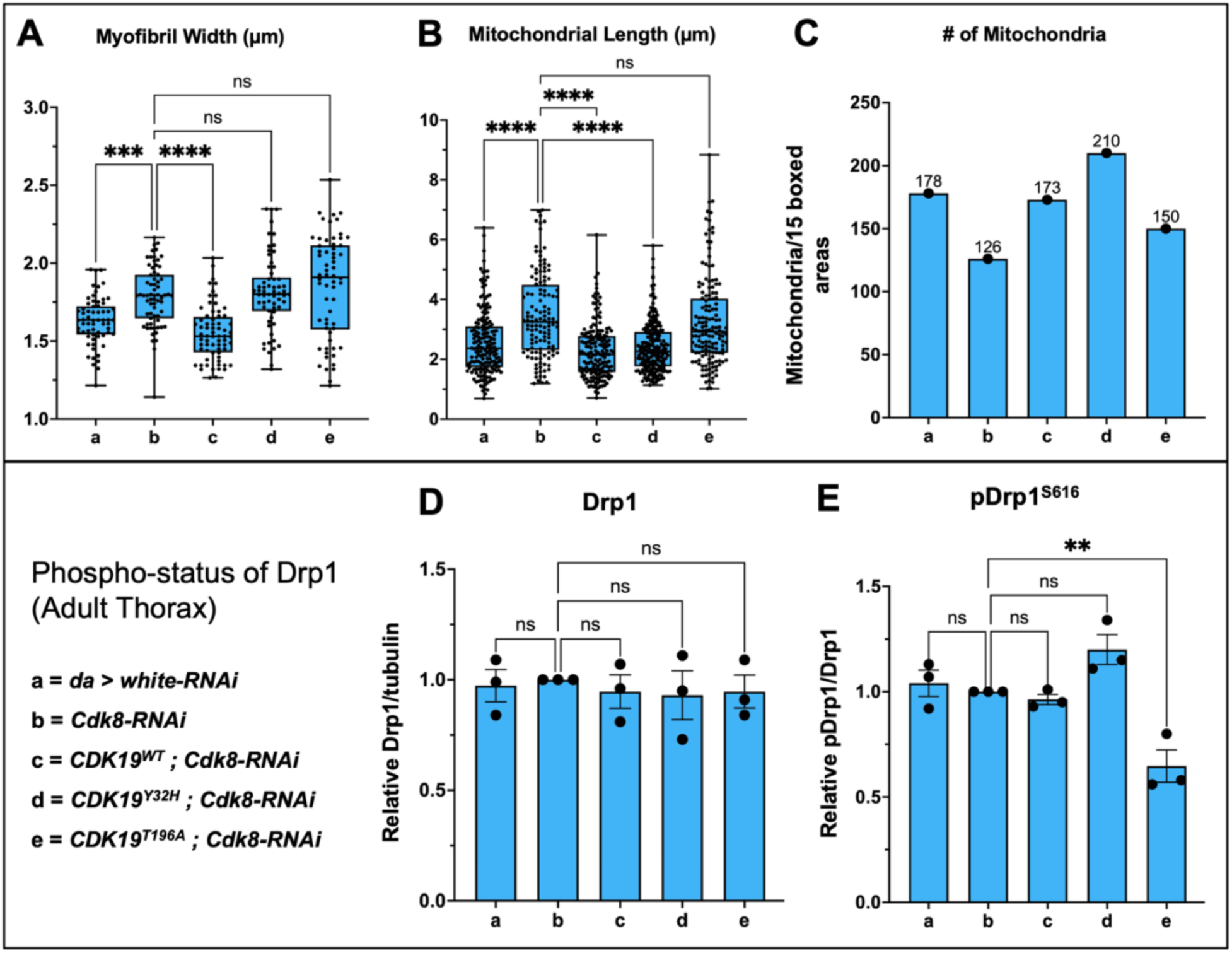
Patient variants of *CDK19, Y32H* and *T196A,* show various effects to muscle, mitochondria, and Drp1 phospho-status compared to *cdk8* knockdown alone. Quantification of **(A)** adult fly indirect flight muscle myofibril width, **(B)** mitochondrial length, with respect to *cdk8-RNAI* and **(C)** mitochondrial counts per 15 assessed regions of muscle. Muscle phenotype data shown as box and whisper plots with solid line at the mean and whiskers for range with 15 muscle regions assessed per genotype. Mitochondrial counts shown as bar plots. **(D)** Relative Drp1/tubulin loading control and **(E)** relative pDrp1/Drp1 from adult thorax lysate samples. Results are presented as bar plots of mean intensity ± s.e.m. All muscle tissue is from 3-day old adult males, raised at 25°C, with n = 5 unique muscle tissue per genotype repeated in n=3 biological replicate immunofluorescent staining or lysate preparations respectively. Statistical analyses are ordinary one-way ANOVA followed by a Dunnett multiple comparisons test. (**p < 0.01 ***p < 0.001, ****p < 0.0001, ns no significance).

**Figure S5.**
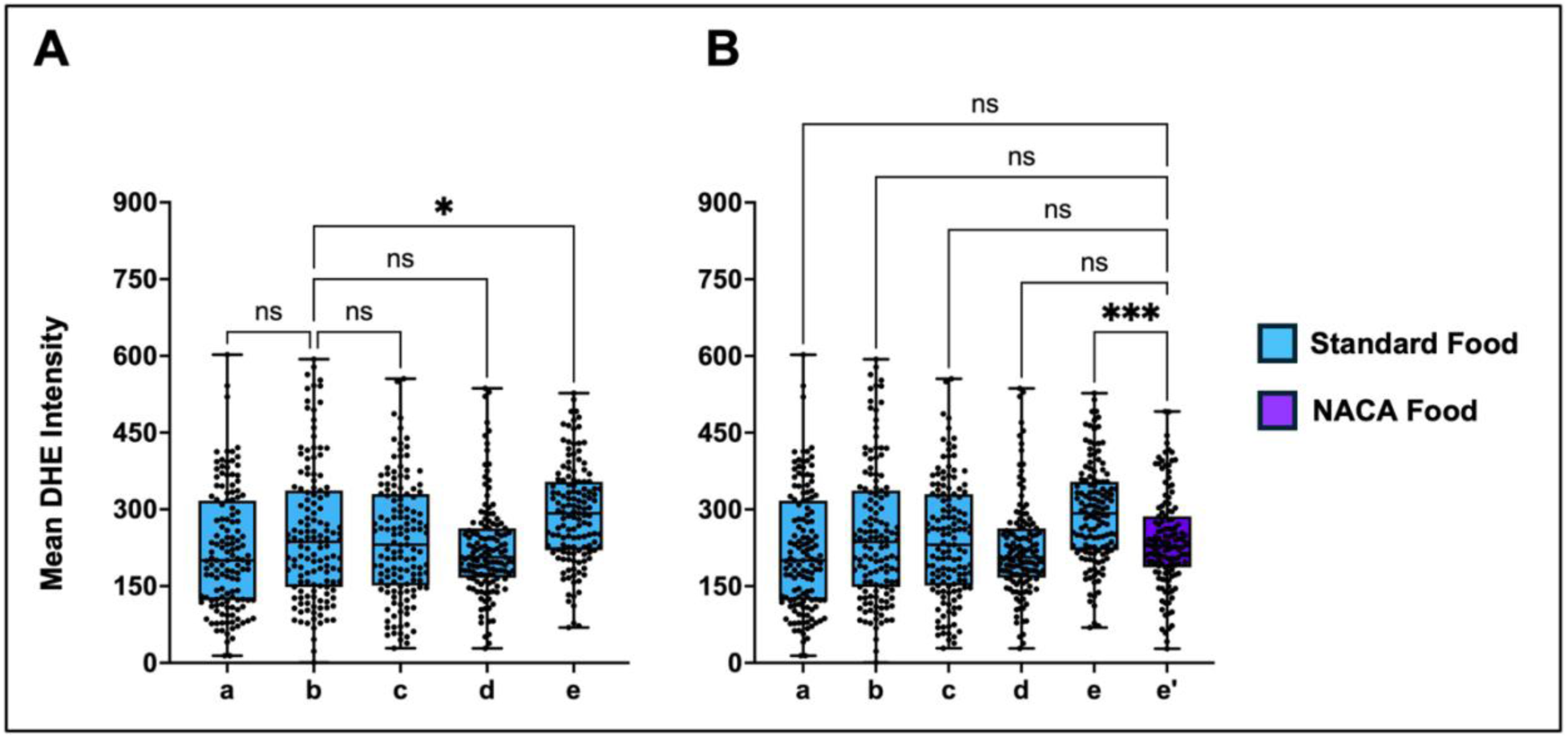
T196A adult muscle ROS is significantly elevated compared to the *cdk8* knockdown group while the Y32H group is unaffected. (A) Quantification of mean DHE intensity from adult fly IFM from standard food raised progenies compared to *cdk8-RNAi*. **(B)** Quantification of mean DHE intensity from adult fly IFM from standard food raised progenies compared to *T196A* group raised on NACA food. Quantifications shown as box and whisper plots with solid line at the mean and whiskers for range with n = 9 muscle regions assessed per genotype. All muscle tissue is from 3-day old adult males, raised at 25°C, with n=9 muscle across three biological replicates. Statistical analyses are ordinary one-way ANOVA followed by a Dunnett multiple comparisons test. (*p < 0.05, ns for no significance).

**Figure S6.**
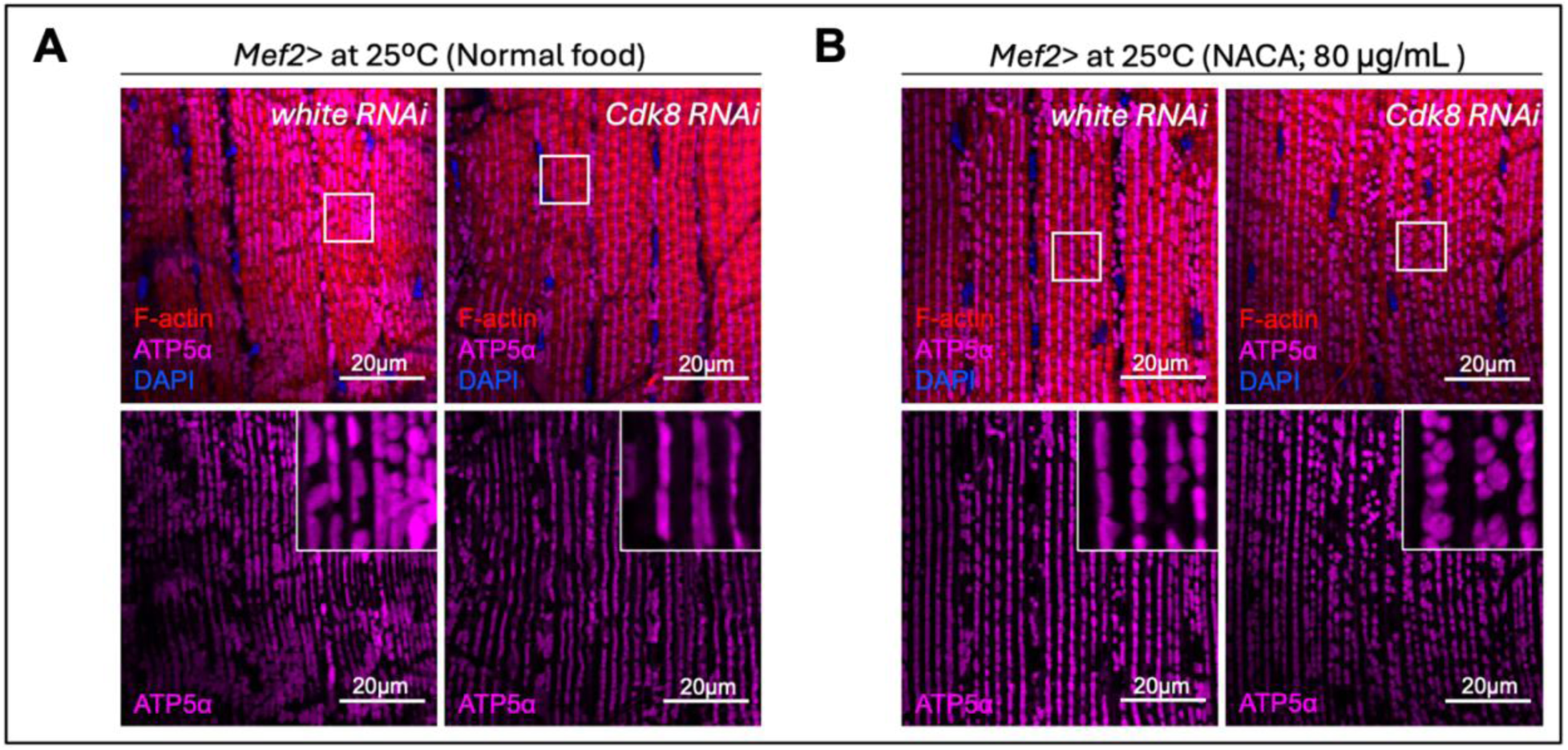
Preliminary assessment of the effects of NACA supplementation on *cdk8-RNAi* adult indirect flight muscle mitochondria compared to *white-RNAi.* Rhodamine-phalloidin staining (F-actin, red), ATP5α staining (mitochondria, green), and DAPI staining (DNA, blue) to visualize adult indirect flight muscle mitochondrial morphology in the indicated genotypes. Scale bar: 5 µm. **(A)** Adult indirect flight muscle of adult flies raised on standard food medium compared to **(B)** NACA food medium.

